# Ten years and a million links: building a global taxonomic library connecting persistent identifiers for names, publications and people

**DOI:** 10.1101/2023.05.29.542697

**Authors:** Roderic D. M. Page

**Affiliations:** School of Biodiversity, One Health and Veterinary Medicine College of Medical, Veterinary and Life Sciences University of Glasgow, Glasgow G12 8QQ

## Abstract

A major gap in the biodiversity knowledge graph is a connection between taxonomic names and the taxonomic literature. While both names and publications often have persistent identifiers (PIDs), such as Life Science Identifiers (LSIDs) or Digital Object Identifiers (DOIs), LSIDs for names are rarely linked to DOIs for publications. This article describes efforts to make those connections across three large taxonomic databases: Index Fungorum, International Plant Names Index (IPNI), and the Index of Organism Names (ION). Over a million names have been matched to DOIs or other persistent identifiers for taxonomic publications. This represents approximately 36% of names for which publication data is available. The mappings between LSIDs and publication PIDs are made available through ChecklistBank. Applications of this mapping are discussed, including a web app to locate the citation of a taxonomic name, and a knowledge graph that uses data on researcher’s ORCID ids to connect taxonomic names and publications to authors of those names.

## Introduction

One thing the field of biodiversity informatics has been very good at is creating databases. However, this success in creation has not been matched by equivalent success in creating deep links between those databases (Thomas, 2009). Instead we create an ever growing number of silos. An obvious route to “silo-breaking” is the shared use of the same persistent identifiers for the same entities across those databases. For example, rather than mint its own identifier for a publication, a database could reuse the existing Digital Object Identifier (DOI) for that publication. This seemingly trivial step of reusing someone else’s identifier opens up numerous possibilities for interconnection, but comes with some risk: what if that persistent identifier does not, in fact, persist? If we cannot trust that an identifier will continue to be maintained and resolve as we expect, then anything we ourselves build upon that identifier is likely to break. Cross linkages between databases are more likely to be made between databases that make efforts to maintain their identifiers (Shorthouse, 2020).

DOIs are a well known example of a persistent identifier, widely used to identify academic publications and other digital items, including datasets. They have been adopted by publishers, who routinely include DOIs for articles from other publishers in the lists of literature cited in their own publications. Embedding these identifiers in PDFs that are intended to be long-lived versions of record requires a significant degree of trust. In particular, the publishers trust that the persistence of these identifiers will be longer than the typical decade lifespan for web links (Hennessey & Ge, 2013). This identifier persistence, coupled with tools to retrieve machine-readable metadata for items with DOIs has led to an ecosystem of services that depend on (or make use of) DOIs, including the citation graph (Heibi et al., 2019), measures of attention (e.g., https://www.altmetric.com/), populating bibliographies for researchers, and machine learning tools to summarise and interpret article content (Nicholson et al., 2021).

DOIs have gained wide acceptance as identifiers of digital publications and data, and have also been adopted for bacterial taxa and their names (Garrity & Lyons, 2003), and for species hypotheses for fungi (Nilsson et al., 2019). However, the bulk of the taxonomic community went a different route and adopted Life Science Identifiers (LSIDs) for taxonomic names. These identifiers were attractive for several reasons: they were developed within the life science community, natively supported the Resource Description Format (RDF), and are free. Core taxonomic databases such as Index Fungorum, International Plant Names Index, and the Index of Organism Names all supported LSIDs, including their novel resolution mechanism. In subsequent years the ability and/or willingness of databases to support LSIDs has declined until few now do so natively (but work-arounds such as HTTP resolution are still feasible, such as https://lsid.io). Despite this, LSIDs are still being embedded in taxonomic publications as part of pipelines to register new taxonomic names (Penev et al., 2016).

An additional problem has been the lack of a single, definitive identifier for the same taxonomic name. New plant names typically have LSIDs issued by IPNI. Due to its origins as a combination of three different databases (Croft et al., 1999), IPNI has duplicate names and hence multiple LSIDs for the same name. Fungal names may have LSIDs issued by Index Fungorum, and URLs issued by Mycobank (Robert et al., 2013). These identifiers share the same local identifier (an integer) and so can be regarded as interchangeable.

In zoology the situation is more complex. Registration of new names is managed by ZooBank, which mints LSIDs for new names, and also has LSIDs for some older names. However the 326,000 records currently in ZooBank represent a small fraction of described animal species. For example, ION has over 5 million names, each with a LSID. Clustering the ION names for duplicates (R. D. Page, 2013) reduces the total to approximately 4.3 million, still considerably more than in ZooBank, or in any other zoological name aggregator such as WoRMS (WoRMS Editorial Board, 2023). The existence of multiple identifiers for the same name complicates attempts to cross-link databases, because it is not obvious which taxonomic name identifier to use. In the absence of a synthesis of these identifiers by the taxonomic community, we may have to rely on third-party identity brokers such as Wikidata (Veen, 2019) to manage cross-links between the menagerie of zoological databases.

### Linked data is not enough

The less than satisfactory history of persistent identifiers for taxonomic names may suggest that the problem was the choice of identifier (e.g., LSID rather than, say, DOI). But this overlooks the deeper problem that, as implemented, LSIDs offered little of value beyond their persistence. Resolving a LSID typically returns RDF with no external links, that is, no identifiers beyond ones local to the LSID provider. We had, in effect, created yet another data silo, ironically using the data format that was supposed to be a silo-breaker.

Currently RDF is enjoying something of a renaissance, especially when serialised as JavaScript Object Notation for Linked Data (JSON-LD) which is more readable and developer friendly than formats such as RDF XML. A growing number of websites relevant to biodiversity are embedding JSON-LD in their pages (see list at https://github.com/rdmpage/wild-json-ld/), including prominent databases such as the Catalogue of Life (https://www.catalogueoflife.org). Yet many of these linked data-enhanced web sites are still silos. For example, the JSON-LD for *Cordyceps changchunensis* from the Catalogue of Life shown in Fig. 1 lacks external identifiers for either the taxonomic name *Cordyceps changchunensis* or the publication of that name. These identifiers exist (urn:lsid:indexfungorum.org:names:839249 and https://doi.org/10.3897/mycokeys.83.72325, respectively). Including them (Fig. 2) converts a data silo into a record connected to two other data sources, and potentially through those sources connected to an even wider network of information.

**Figure 1.**
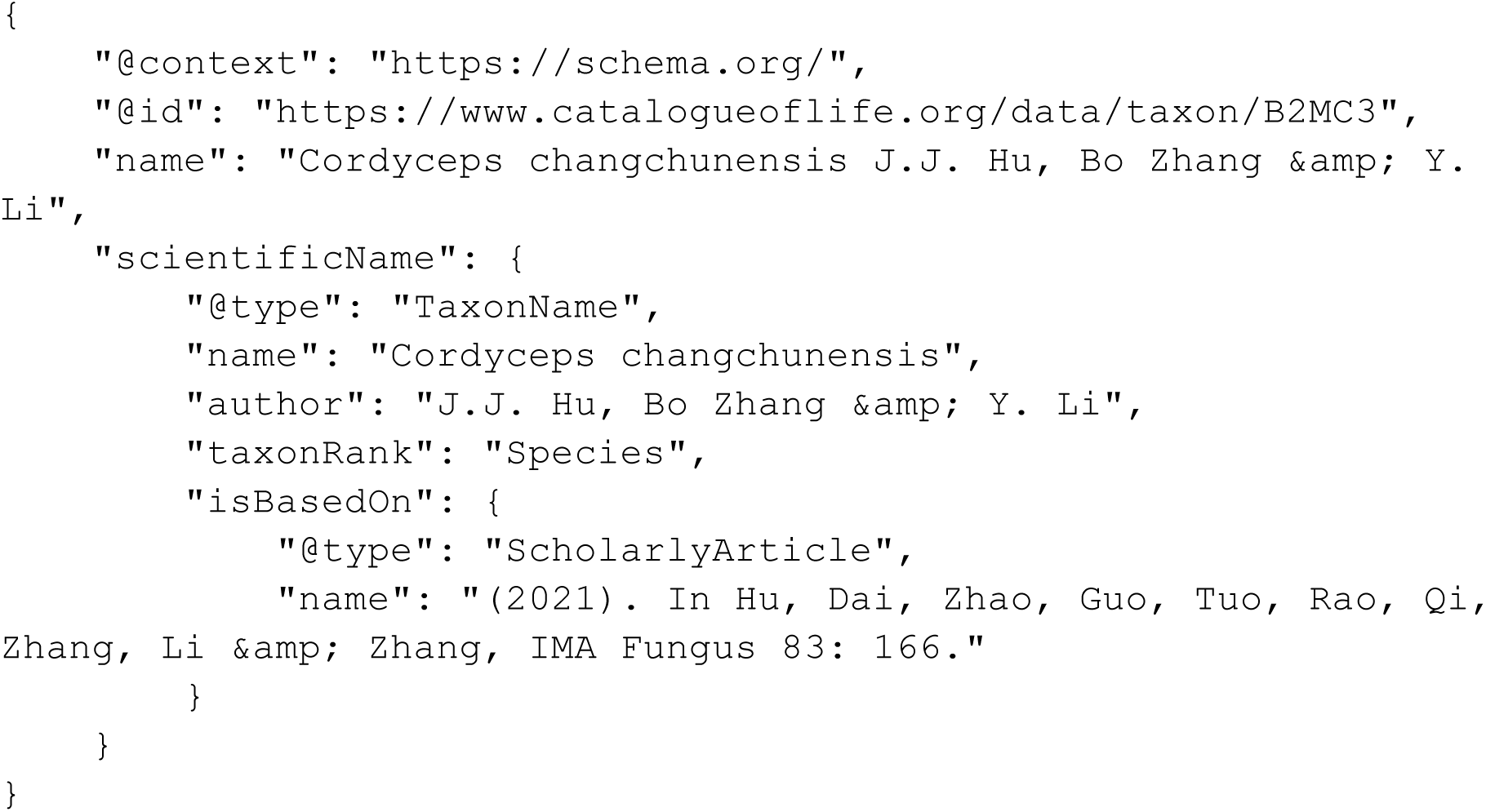
Simplified JSON-LD for Catalogue of Life taxon B2MC3, retrieved 16 May 2023. Note the lack of an identifier for either the scientific name or the publication that name is based on.

**Figure 2.**
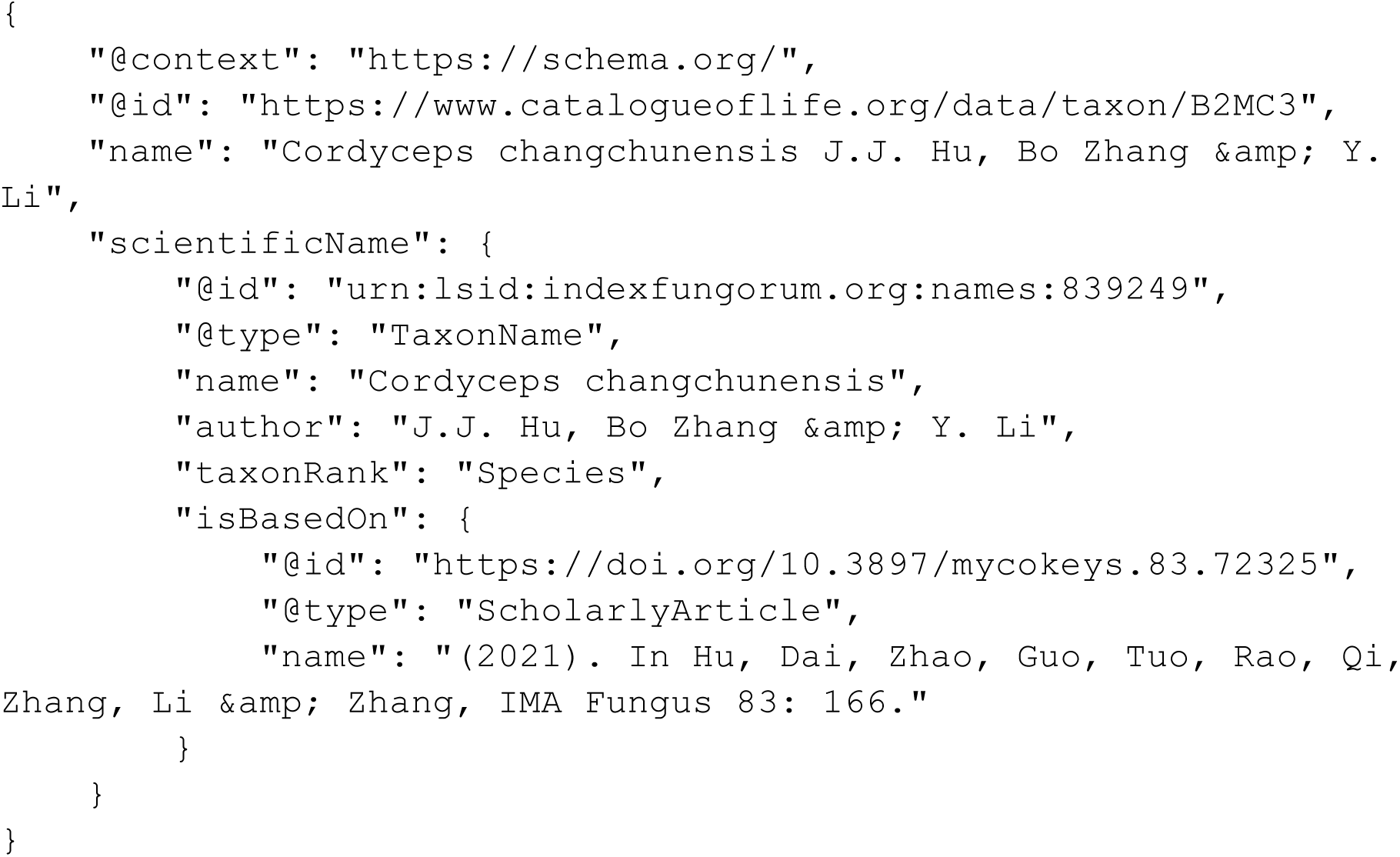
The JSON-LD shown in Fig. 1 enhanced by including persistent identifiers for the taxon name (LSID) and its publication (DOI).

### Putting holes in silos

The goal of the work described here is to make a small hole in taxonomic data silos by linking LSIDs for taxonomic names to DOIs for the works that published those names. Other bibliographic identifiers are also available and relevant, but the focus in this work will be on DOIs. One reason for this focus is that DOIs are a relatively “sticky” identifier that is likely to be connected to other identifiers, most notably ORCID ids for researchers (Bohannon & Doran, 2017). Another reason is the role DOIs play in creating the citation graph, the scholarly network linking works to the works that they either cite or are cited by (Shotton, 2013). It also makes it easier to cite the taxonomic literature. Taxonomists frequently complain about the lack of citations their work receives. Whatever the merits of that complaint, calls for better citation practices (Benichou et al., 2022) are unlikely to improve the situation if the taxonomic literature remains disconnected from taxonomic names. How are we to know what publications should be cited for a name if the links between names and literature are hard to discover?

### Storing the mapping

In addition to the challenge of creating these mappings, there is the problem of how to make them available for reuse. Ideally the source taxonomic databases would incorporate them, on the grounds that they would add value to their users, and it would save those databases doing the work themselves. However, this assumes that those databases are willing, or have the resources to incorporate this additional data, which rarely seems to be the case. Alternative approaches include developing separate, stand alone web sites to make the data available, or simply putting a data dump in a repository.

I have experimented with various approaches. In 2013 I created a standalone database mapping ION LSIDs to DOIs and other identifiers, and wrapped this in a user-friendly web site (https://bionames.org) developed with funding from the Encyclopaedia of Life (R. D. Page, 2013). In 2018 I explored an intermediate approach of using Datasettes to publish a mapping between IPNI names and the literature (R. Page, 2018). This made the data available and queryable, but the interface doesn’t support taxonomic-specific queries. Both these approaches result in standalone web sites with little obvious means to integrate the mapping into other databases.

The recent release of ChecklistBank (https://www.checklistbank.org) (Döring et al., 2022) has provided a new way to publish the data so that it complements existing databases. ChecklistBank includes all the taxonomic checklists used to create the Catalogue of Life, but also enables users to upload their own checklists. This means that we can take a taxonomic checklist, add persistent identifiers for the literature, then upload the augmented data to ChecklistBank as a new dataset (with an appropriate citation to the original source database). This augmented checklist can have its own DOI and be citable (hence providing a mechanism to give credit to those making the links). Because the augmented dataset uses the same taxon name identifiers as the original database, this also means that at any point the original data publishers could incorporate literature mapping into their own databases. Likewise, any other database that uses those same taxon name identifiers could also use the mapping.

ChecklistBank provides a convenient way to store mappings between names and publications, but this is a single edge in the biodiversity knowledge graph (R. Page, 2016). Storing the deeper links, such as between taxonomic names, publication, people, institutions, and funders requires more flexibility. To store these I follow an approach sketched in (R. Page, 2022) where the mappings are stored as RDF and published to Zenodo. These mappings can then be loaded into a triple store.

### Goals

The goal of this work is to make available over a million links between persistent identifiers for taxonomic names and the publications for those names. This paper covers three databases: Index Fungorum (https://www.indexfungorum.org), IPNI (https://www.ipni.org), and ION (http://www.organismnames.com). Each of which uses LSIDs as persistent identifiers for taxonomic names. This gives us substantial coverage of animals, plants, and fungi.

In this work I will focus on "citable" bibliographic identifiers, that is, identifiers that are typically cited by other publications. In practical terms this means DOIs (Fig. 3). The two advantages of work-level identifiers is that they do tend to be persistent (e.g., DOIs) and they are also the basis of measures of scientific activity (e.g., citations) and attention (e.g., altmetrics).

**Figure 3.**
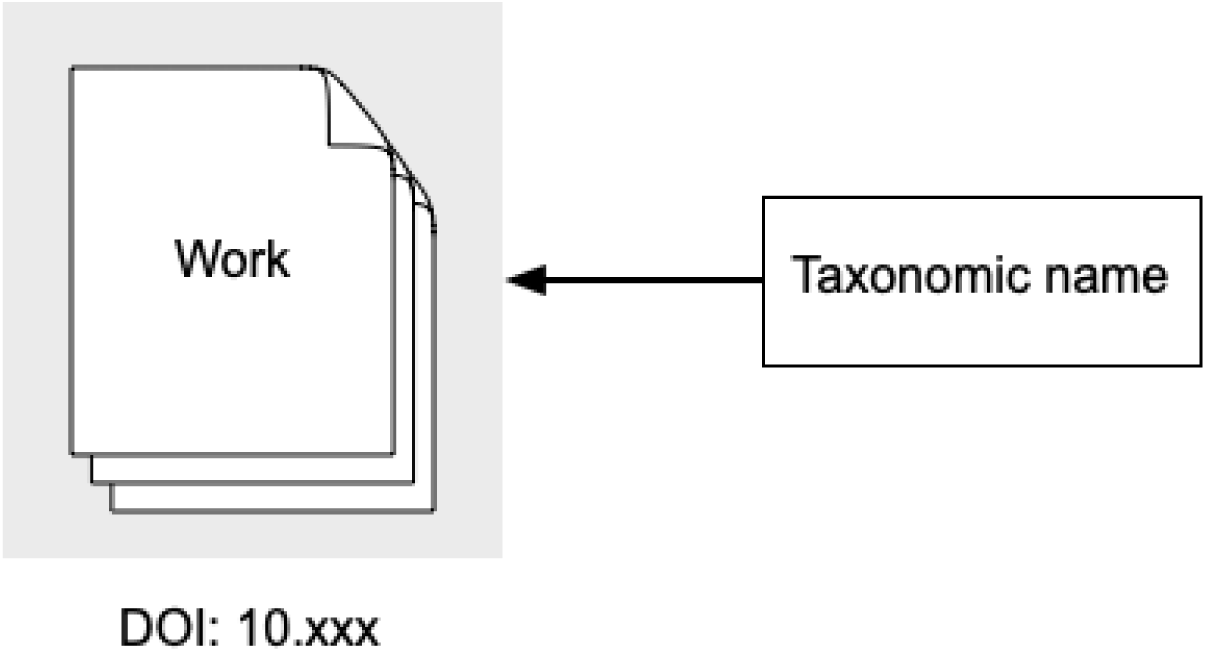
Work-level identifiers. The taxonomic name is linked to a work-level identifier, such as the DOI for the article that published the name.

In contrast, databases such as IPNI and IF typically store bibliographic information at the level of individual pages, or sets of pages. Citations at the page level have been termed "microcitations" and are analogous to what the U.S. legal profession refers to as "point citations" or "pincites". Some bibliographic databases support page-level identifiers. For example, individual pages in the Biodiversity Heritage Library (BHL) have their own unique URL. In cases where there isn’t an explicit identifier we can use “fragment identifiers” to identify parts of an entity (Fig. 4). For instance, an individual page in a PDF can be referred to using the fragment #page=*n* where *n* is the position of the page within the PDF, starting from *n*=1 for the first page (Taft et al., 2004). Blocks of text within a page can be identified using TextQuotes or TextPosition identifiers (Dürst & Wilde, 2008). Locations with a HTML or XML document can be referred to using XPath statements. Fragment identifiers enable deep within-document linking, but can be fragile. If the document being linked to changes, or has multiple versions, then fragment identifiers may no longer successfully link to the desired content (Brush et al., 2001).

**Figure 4.**
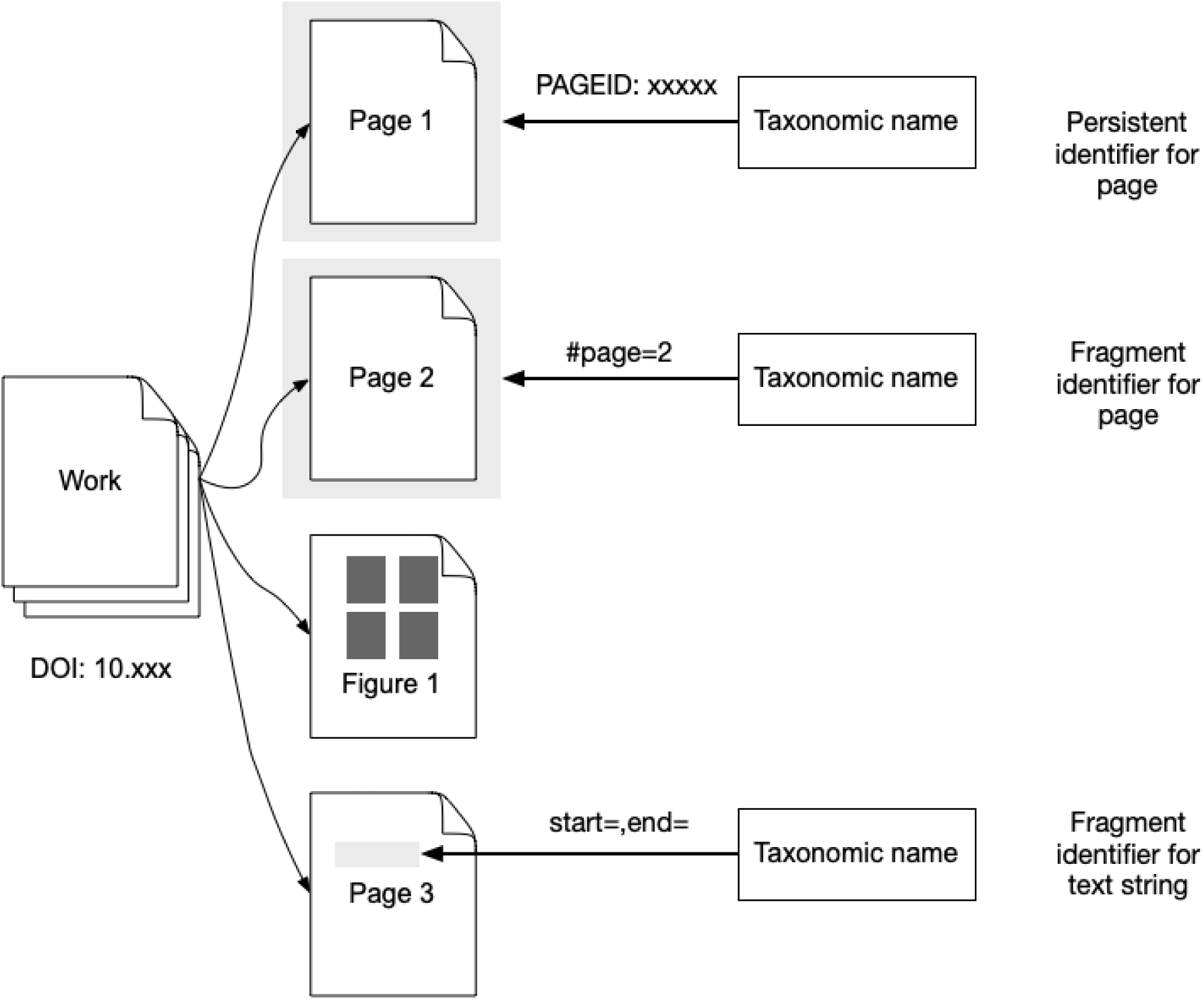
Page and fragment-level identifiers. In contrast to work-level identifiers (Fig. 3) we can use identifiers for pages or parts of pages.

Another approach is to select one or more blocks of text and any associated figures within a publication and treat that collection as a distinct unit (Fig. 5). These can be treated as stand alone entities, or recombined to provide an alternative navigation pathway through a set of papers (‘Spinning Threads’, 2012). The Plazi project (Agosti & Egloff, 2009) extracts blocks of text and images as “treatments”, and many of these are assigned DOIs. This has the advantage of creating citable units, although to date there is little evidence that either taxonomic databases or publications actually cite treatments rather than the entire work.

**Figure 5.**
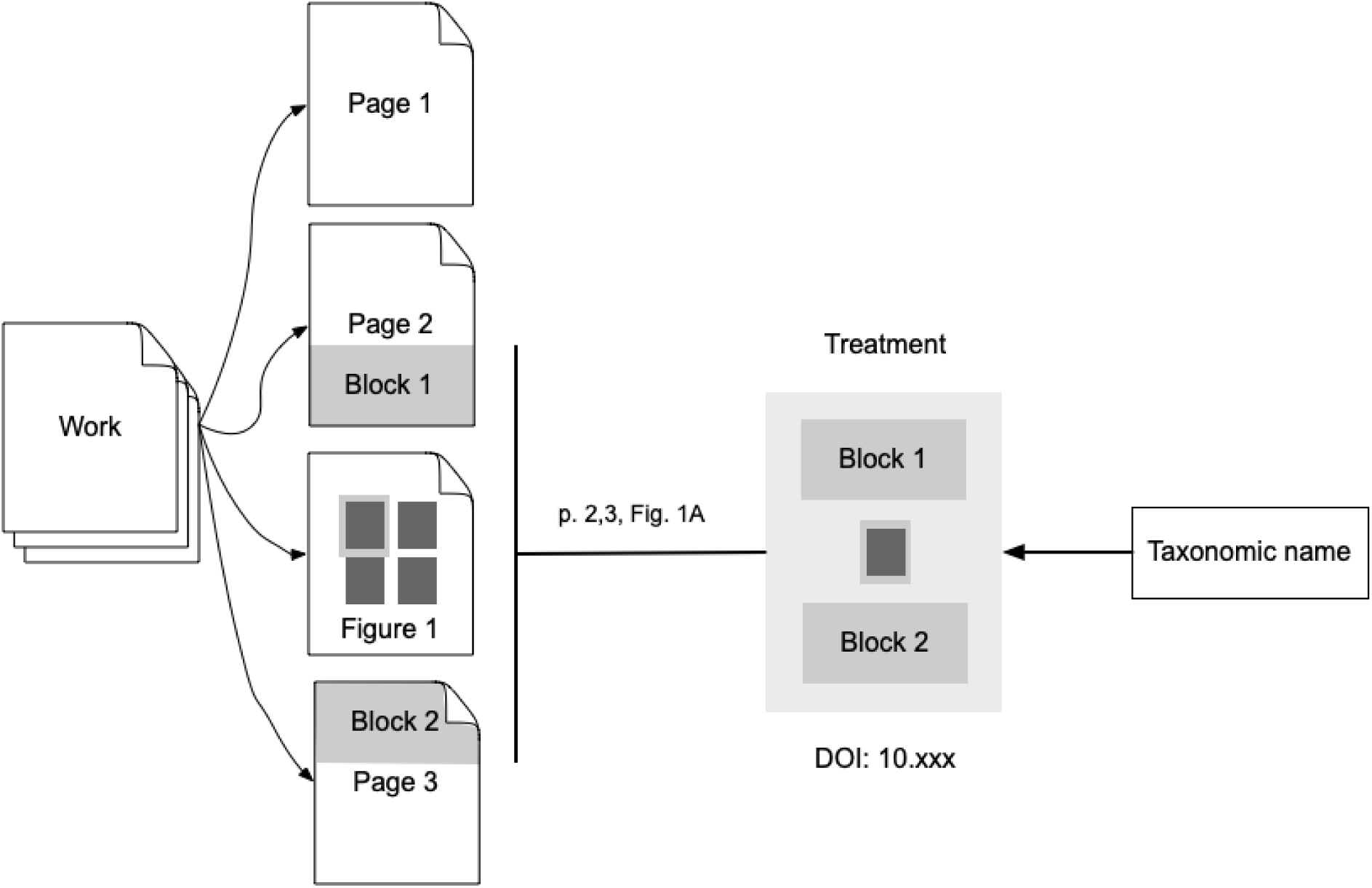
Part of work identifiers. Taxonomic names are linked to one or more parts of a document, such as text and a figure. These parts are packaged into a citable unit such as a treatment.

### Outputs

There are three categories of output from this work. The first is a mapping between LSID for a taxon name and one or more (ideally) persistent identifiers for the publication that established that name. Where available a DOI is used, but other identifiers are also included, such as Handles and URLs. If the publication has an item in Wikidata, the QID of the Wikidata item is also included. These mappings are published to ChecklistBank.

The second category of output is a version of the mapping in RDF, enabling the mapping to be used in a knowledge graph. In this paper I combine the mapping with data from ORCID sufficient to construct a simple knowledge graph of taxonomic names and the taxonomists who published those names.

The final category of output is a proof-of-concept web site that uses the mappings stored in ChecklistBank to generate citations for publications of a taxonomic name, as well as locate a PDF for that article.

## Methods

### Taxonomic names

Names and citation data were obtained from IF, IPNI, and ION at various times over the last decade. Typically data was retrieved by resolving LSIDs for individual names, parsing the resulting RDF into tabular form and storing this data in SQL databases for ease of manipulation. If the database no longer supports LSID resolution, tools such as https://lsid.io can be used to retrieve the RDF. On other occasions bulk downloads have been made using APIs provided by the databases. Once in a local SQL database the data has been cleaned, citation strings parsed, any existing bibliographic identifiers extracted, then the citation data is mapped to external bibliographic identifiers.

### Mapping citations

For full citations that include data such as authors, title, journal, and pagination, there are a number of approaches to mapping these citations to identifiers. These include search engines such as CrossRef or ReFindit (https://refindit.org/about.html). Most tools have their own unique search interface, but some support generic search interfaces such as the Open Refine API.

Matching full citations can be treated as a simple string matching task. However, microcitations (or “pincites”) present an additional challenge. The simplest micro citation is a single page within a publication. If we have a database of page ranges for articles (i.e., the start and end page numbers) then matching microcitations to full citations is relatively trivial: find the article in a given volume that has a page range that includes the page in the microcitation. However, given that we lack a freely accessible database of all taxonomic publications this can be a challenge. It also assumes that available metadata for articles includes page numbers. In some cases these numbers are not readily available, for example for the *European Journal of Taxonomy* of 1,201 articles from 2011 - 2023 only 243 had a page range in the CrossRef metadata. Another reason for the lack of page numbers is the move to online publication where the notion of a “page” becomes problematic. Pagination depends on how the article is rendered, and may vary across different representations, or be absent altogether.

To facilitate resolving microcitations I have, for the last decade or more, been building a bibliographic database that includes pagination data. This data comes from a variety of sources, such as CrossRef, PubMed, JSTOR, journal websites, article PDFs, etc. Managing this data locally is essential as often the metadata available from individual sources is incomplete (for example may lack page numbers) and hence multiple sources may be required to retrieve sufficient metadata to determine the appropriate persistent identifier for a publication record in a taxonomic database. To make this data more widely available I am uploading much of it to Wikidata, where it can be further curated and improved (R. D. M. Page, 2022).

Other approaches for mapping citations include using identifiers for articles, or parts thereof, which have also been incorporated into taxonomic databases. For example, the record for the taxonomic name *Neodeightonia mucosa* (urn:lsid:indexfungorum.org:names:840943) cites “Frontiers in Microbiology, volume 12, issue no. 737541”. This corresponds to the DOI 10.3389/fmicb.2021.737541 (note the shared “737541”). This is an argument against the use of “opaque identifiers”. Providing one is aware that information in an identifier might be misinterpreted, non-opaque identifiers (typically based on metadata for the entity being identified) can be a useful aid to making connections between databases. This can be particularly useful in cases where a journal has moved from sequential pagination within a volume to continuous article publication such that every article starts on page 1 (Anonymous, 2014).

One unintended consequence of attempting to map citations is it can expose errors in the taxonomic databases. A mismatch between journal and volume numbers is often a clue that a record is in error. For example, the citation for urn:lsid:indexfungorum.org:names:839249 is “Hu, Dai, Zhao, Guo, Tuo, Rao, Qi, Zhang, Li & Zhang, IMA Fungus 83: 166 (2021)”. There is no such volume for IMA Fungus, however the volume and page number match an article in *MycoKeys* (Hu et al., 2021). Databases inevitably benefit from scrutiny, and making links between databases generates a lot of scrutiny.

### Data management

The mapping between names and publications is managed in a local SQL database, either SQLite or MySQL. A range of custom scripts manages data import and cleaning, and matching bibliographic citations to persistent identifiers. There are also tools to visualise progress, discover gaps, and drill down by taxonomic name, publication, or date. Each mapping project is managed in one or more GitHub repositories.

### Storing the mapping in ChecklistBank

For each database a new entry was created in ChecklistBank. A data release in the Catalogue of Life Data Package (CoLDP) format (Döring & Ower, 2019) was created and uploaded to Zenodo where it received a DOI. The same data is then uploaded to ChecklistBank.

The CoLDP format requires a unique identifier for each bibliographic reference. This was generated using a trigger in the SQLite database. If the reference had a Wikidata QID then that value served as the identifier. In the absence of a Wikidata identifier a new identifier would be generated from one of the persistent identifiers added in the mapping, such as the DOI.

The CoLDP expects a bibliographic citation string. LSIDs for the ION database supply a citation string, but in the case of IF and IPNI complete citations are not available in the original databases as these databases use microcitations. Hence in this case complete citations were generated using tools based on the CSL-JSON format. Bibliographic metadata in this format was retrieved from Wikidata, or via content-negotiation from https://doi.org, then formatted for display.

### Outputting the mapping as RDF

In addition to the COLDP format for ChecklistBank I created linked data files for the names using the N-Triples format. The names were modelled following the draft Bioschemas proposal for taxon names (https://bioschemas.org/TaxonName/), which is followed by the Catalogue of Life. Rather than output the entire mapping as RDF I included only those names that have a publication with a DOI. This is because the DOI is likely to be the only bibliographic identifier found in other datasets that we could potentially link to, such as ORCID. For names that have DOIs for their publications the LSID for that name is linked to the DOI using the schema.org property “isBasedOn” (Fig. 6). For each of IF, ION, and IPNI, the list of N-Triples was uploaded to Zenodo.

**Figure 6.**
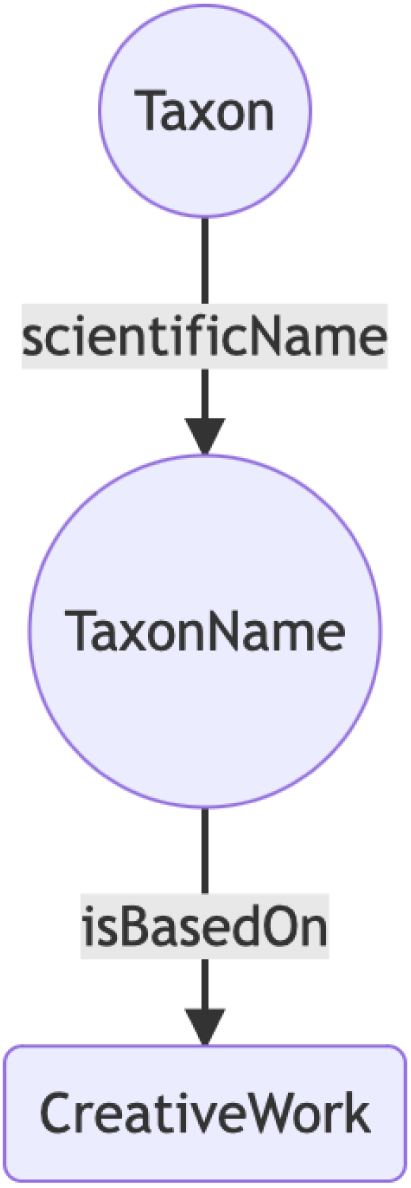
A taxonomic name linked to the publication that makes that name available, expressed using terms from the http://schema.org vocabulary.

### Augmenting with ORCIDs

A knowledge graph is only as interesting as its connections, so to augment the simple pairs of taxonomic names and publications I created an additional RDF file connecting people and their publications. Many researchers have ORCID ids which enables those researchers to uniquely identify themselves (Bohannon & Doran, 2017). The ORCID record for an individual may list their publications (and other outputs, such as data sets, peer reviews, etc.), many of which (but not all) have DOIs. It may also link people to other entities such as organisations where they have studied or worked, or funding agencies (Fig. 7).

**Figure 7.**
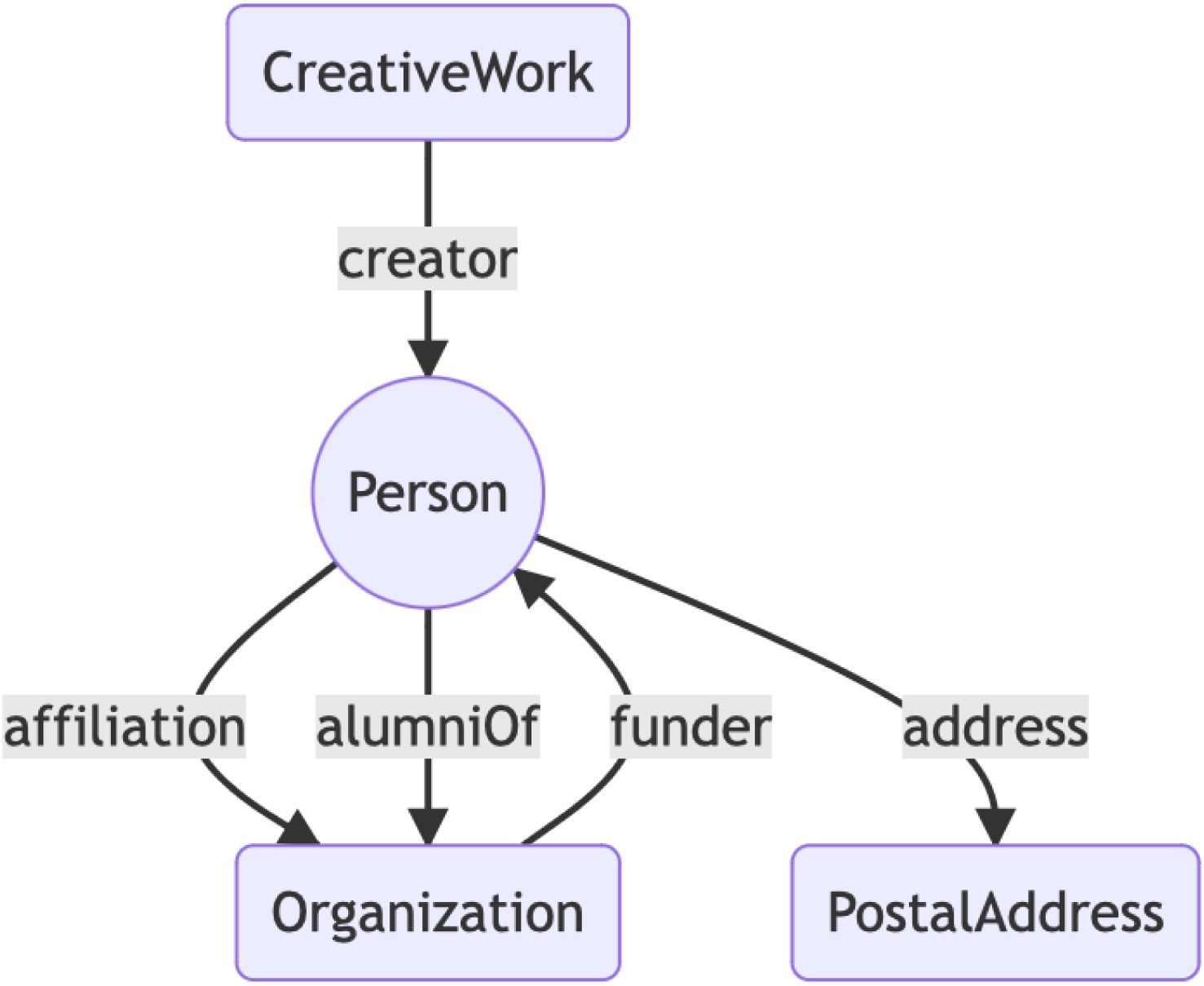
Simplified version of the data model used by ORCID to export data in RDF.

Data in ORCID is available as linked data using content-negotiation. That is, by sending a HTTP request that accepts data in the format “application/ld+json” ORCID will return structured data about a person. I retrieved data for a set of ORCIDs associated with DOis for papers on taxonomy using a tool (https://enchanting-bongo.glitch.me) which queries the ORCID API for ORCID ids associated with DOIs. I also retrieved ORCID ids via queries to Wikidata, for example for authors in Wikidata that have both an ORCID id and an article on Wikispecies).

Much of the data in ORCID is user-supplied, and some of it is messy. A common problem is URLs that aren’t properly formed. Linked data is a very unforgiving format when it comes to URLs, and these errors cause problems when uploading data to a triple store. Hence data from ORCID data was run through a series of scripts to clean extraneous characters and eventually output clean RDF in N-Triples format, suitable for upload into a triple store.

Combining the taxonomic names, literature, and ORCIDs yields a small knowledge graph (Fig. 8).

**Figure 8.**
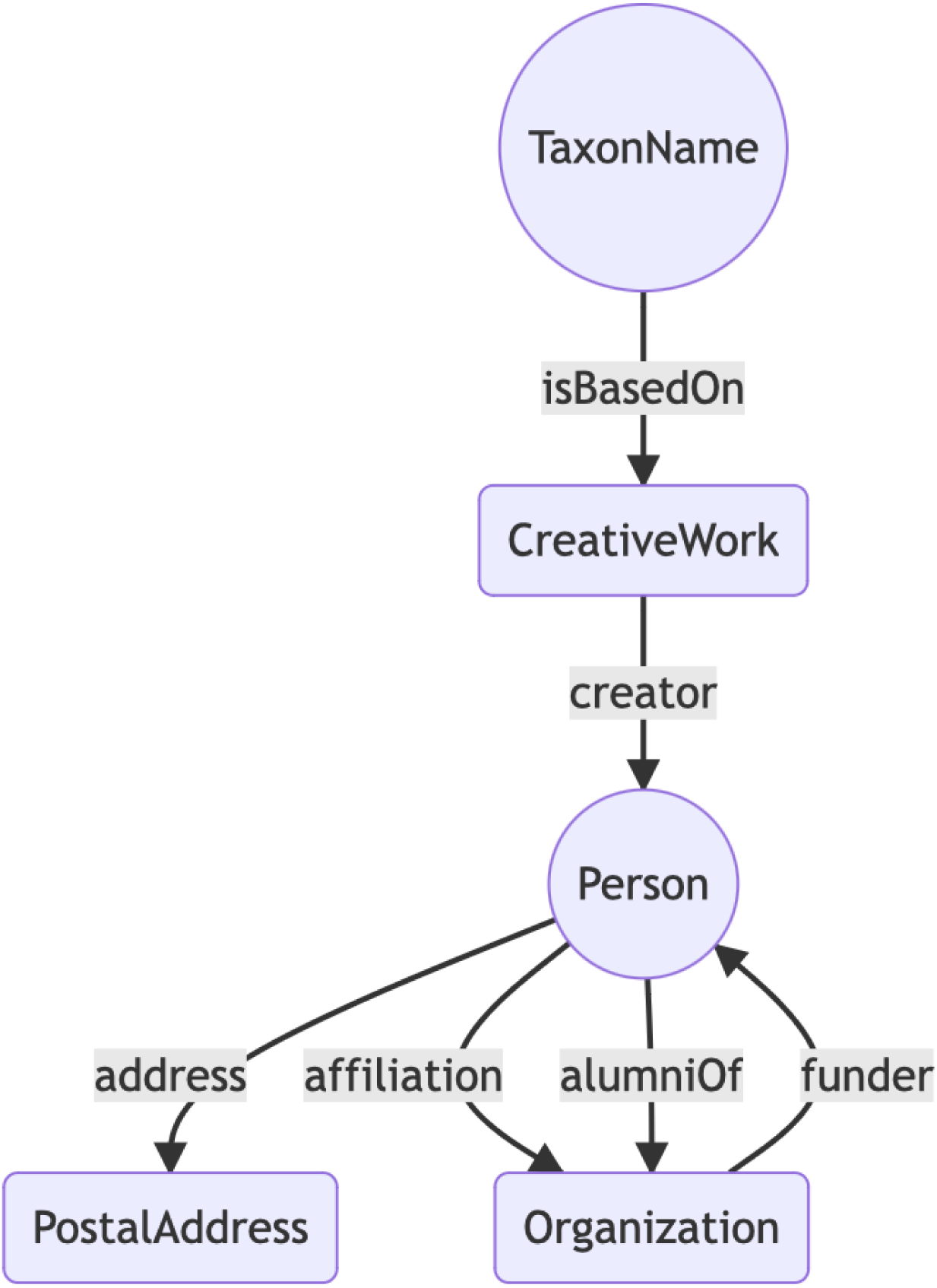
Modelling the relationship between a taxonomic name, its publication, and the author of that publication.

## Results

### Coverage

Table 1 Numbers of taxonomic names and persistent identifiers for publications. For each database the table shows the total number of taxonomic names, how many of those names have publication information, and how many of those publications have been mapped to one or more persistent identifiers. The row “Any” records the number of publications that have any identifier.

**Table 1.**
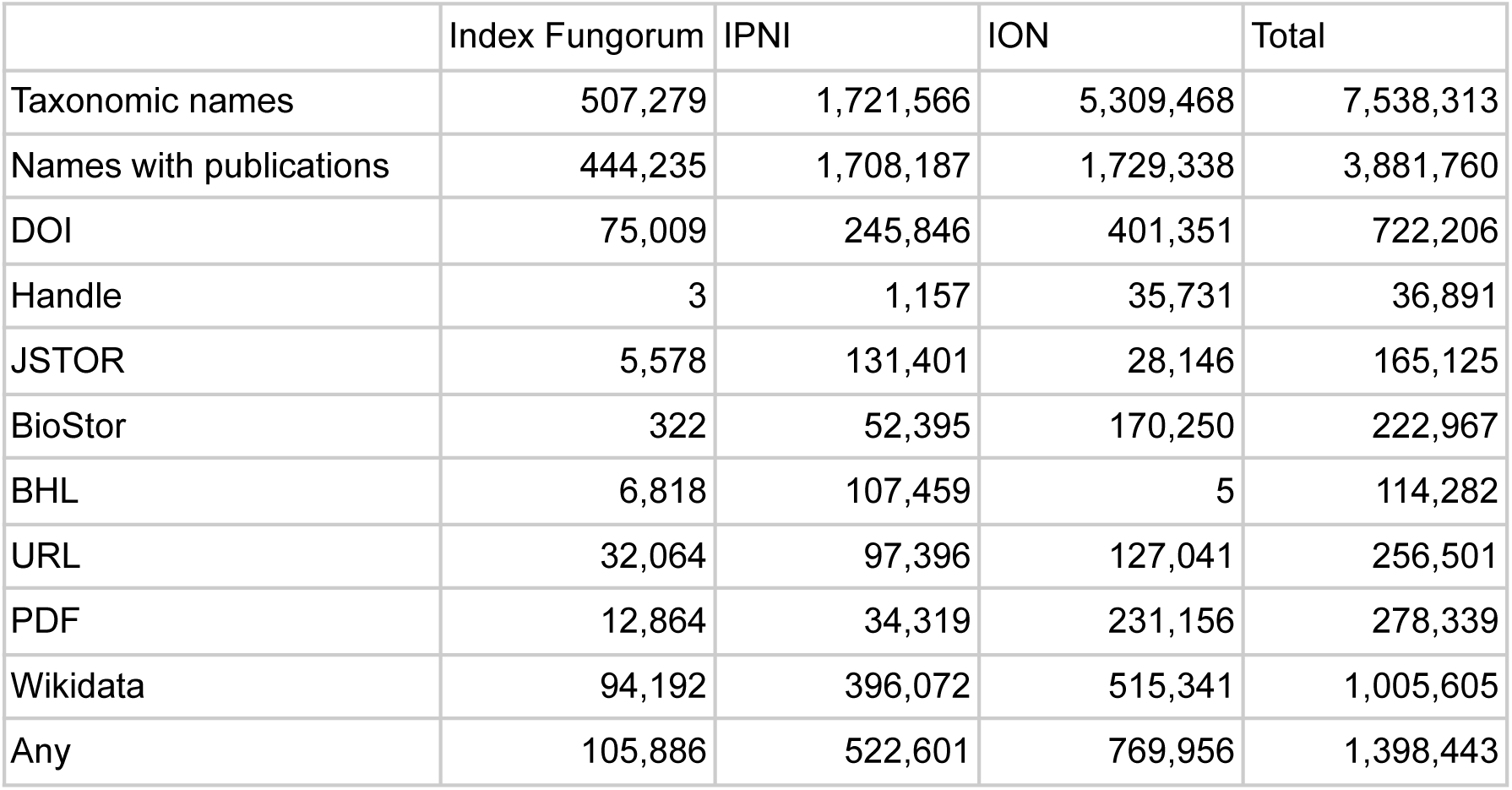
gives basic data on how many names each source database has, and how many have been mapped to a persistent identifier for a publication. The number of names with DOIs for the corresponding publication ranges from 75,000 to 722,000 across the three datasets. Not surprisingly, Wikidata is the single most common identifier, with a little over a million names linked to a publication with a Wikidata QID.

|  | Index Fungorum | IPNI | ION | Total |
| --- | --- | --- | --- | --- |
| Taxonomic names | 507,279 | 1,721,566 | 5,309,468 | 7,538,313 |
| Names with publications | 444,235 | 1,708,187 | 1,729,338 | 3,881,760 |
| DOI | 75,009 | 245,846 | 401,351 | 722,206 |
| Handle | 3 | 1,157 | 35,731 | 36,891 |
| JSTOR | 5,578 | 131,401 | 28,146 | 165,125 |
| BioStor | 322 | 52,395 | 170,250 | 222,967 |
| BHL | 6,818 | 107,459 | 5 | 114,282 |
| URL | 32,064 | 97,396 | 127,041 | 256,501 |
| PDF | 12,864 | 34,319 | 231,156 | 278,339 |
| Wikidata | 94,192 | 396,072 | 515,341 | 1,005,605 |
| Any | 105,886 | 522,601 | 769,956 | 1,398,443 |

To visualise progress on linking names to literature I computed the number of names per decade from 1750 to 2020 that were published in 50 publication venues or “containers” (such as journals, monographs, and books) that published the most names. The publications were then sorted by the decade in which they published the most names (their “modal” decade), which enables us to see changes in the fate of publications over time. The diagrams also plot the percentage of works that have any persistent identifier (or similar, such as a URL).

Both fungi (Fig. 9) and plants (Fig. 10) show similar patterns of apparent turnover in publications, and the most recent publications have the greater density of persistent identifiers. The diagram for animals (Fig. 11) is truncated relative to the other taxonomic groups with no data prior to the 1860’s. This is because Zoological Record, the primary source for ION, only started in 1864 (Bridson, 1968). Many of the top animal publications are still being published, and many have DOIs, which is reflected in the greater density of PIDs in Fig. 11 compared with Fig. 9 and Fig. 10.

**Figure 9.**
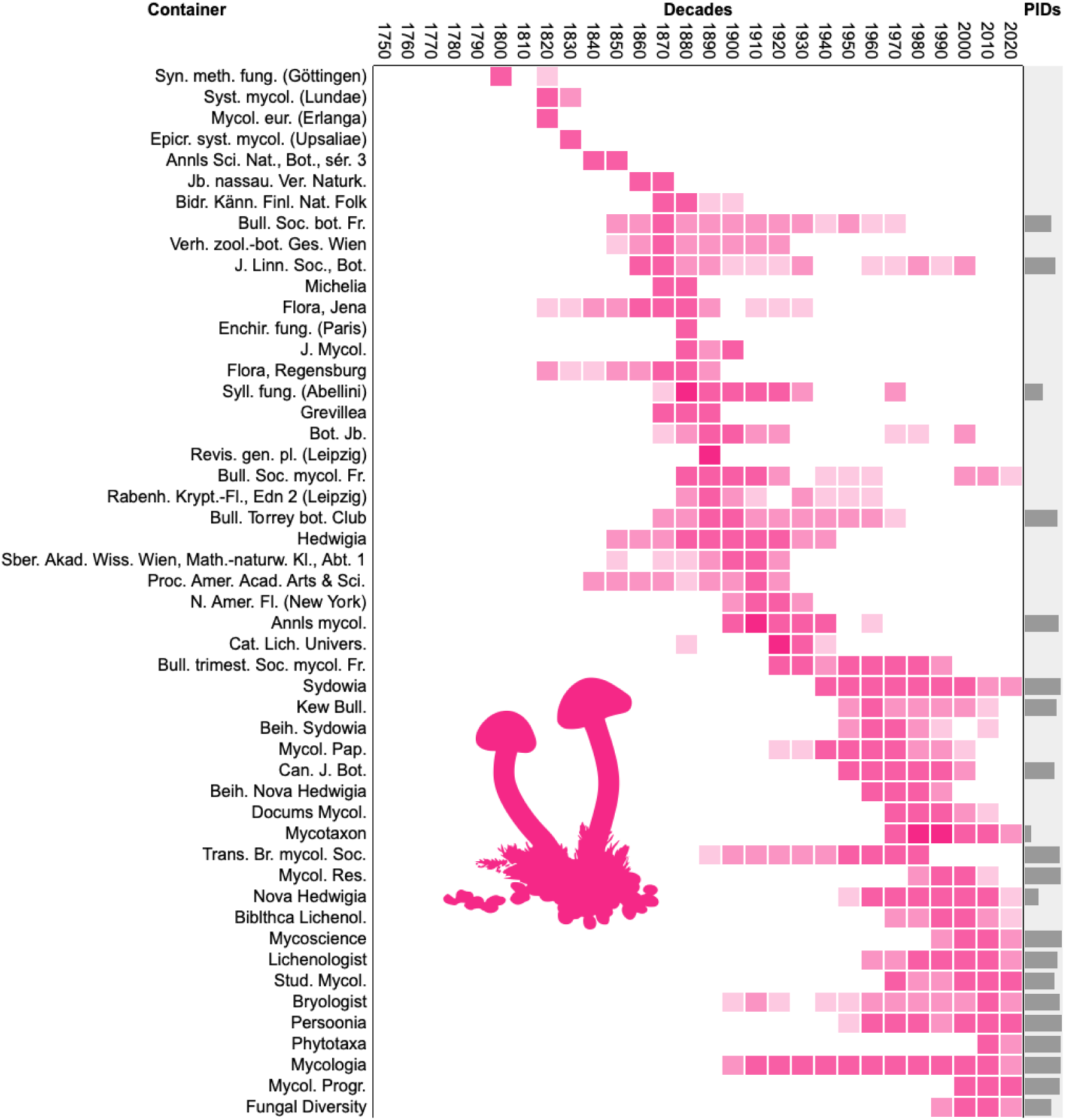
Density distribution of taxonomic names published in the decades from 1750 to 2020 across the 50 publication venues (“containers”) that published the most names for fungi. The containers are ordered by the decade with the largest number of names. The column labelled “PIDs” shows bars proportional to the percentage of names in each publication that have been linked to a persistent identifier for that publication.

**Figure 10.**
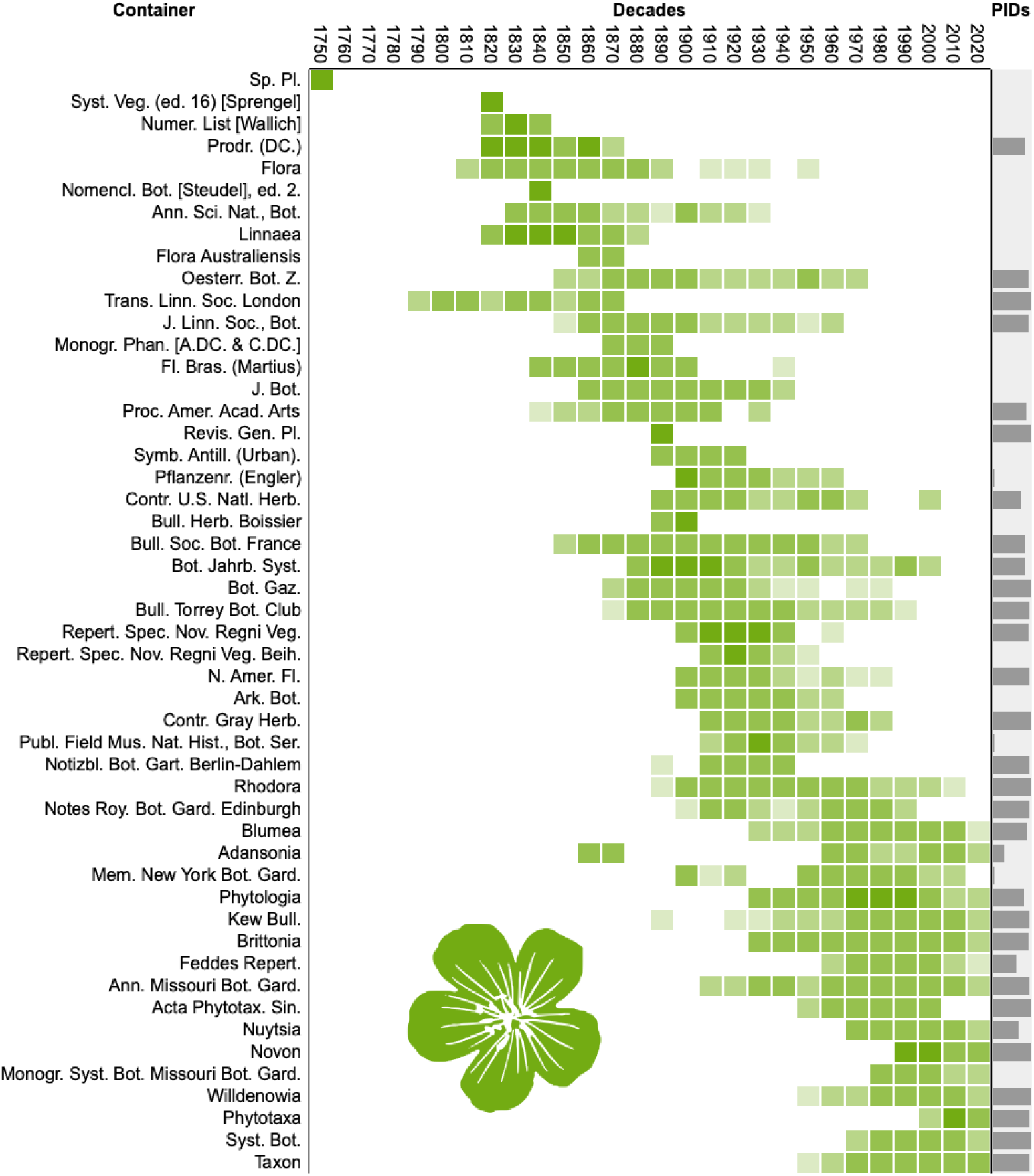
Density distribution of taxonomic names published in the decades from 1750 to 2020 across the 50 publication venues (“containers”) that published the most names for plants. See Fig. 9 for further explanation.

**Figure 11.**
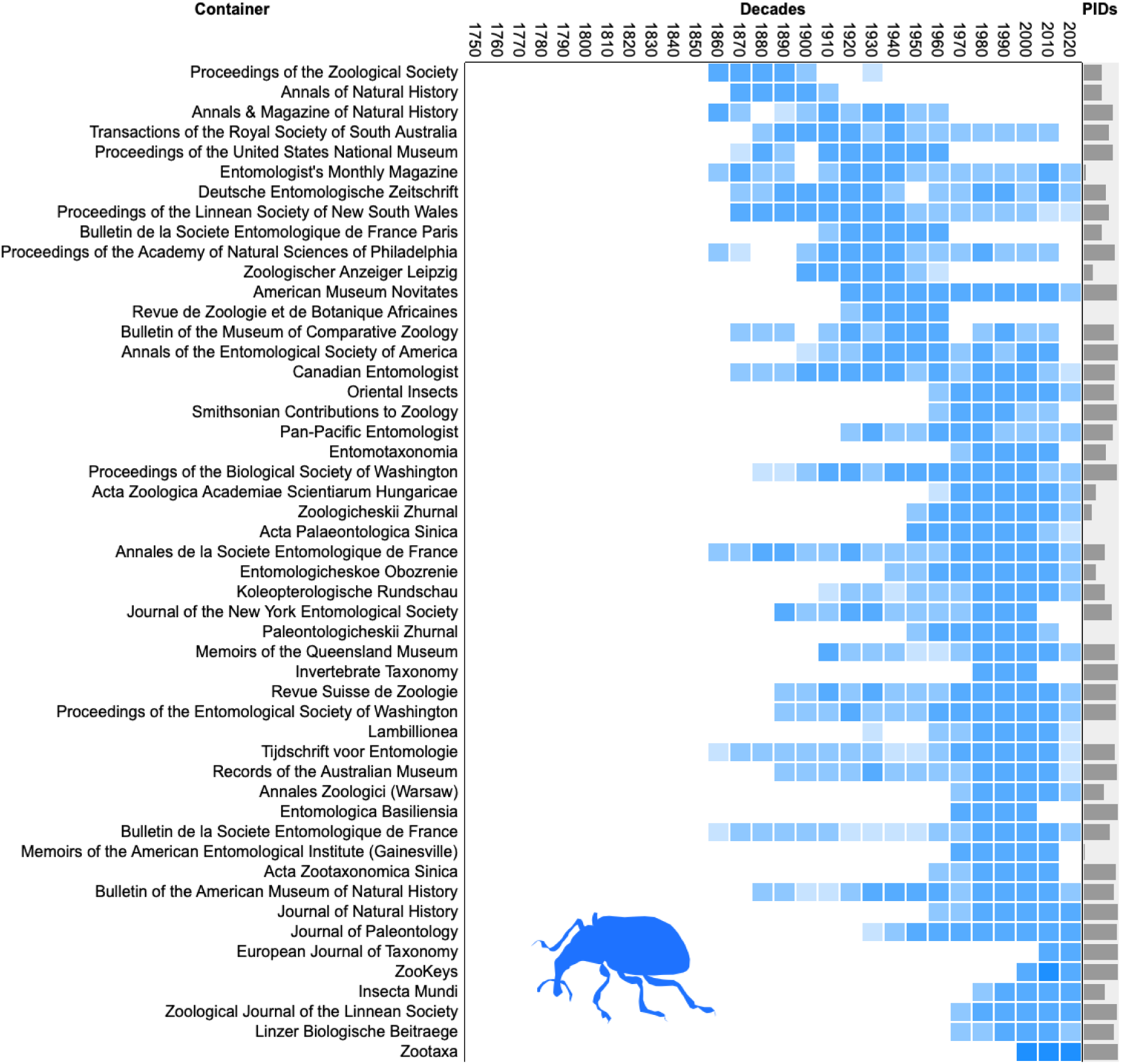
Density distribution of taxonomic names published in the decades from 1750 to 2020 across the 50 publication venues (“containers”) that published the most names for animals. See Fig. 9 for further explanation.

Further differences between fungal, plant, and animal taxonomic publishing can be seen when we plot the total number of names published in each of the top 100 publications (Fig. 12). Animal names are dominated by the journal Zootaxa which has published five times as many names as the next publication (ZooKeys). The size of publication rapidly drops away such that the top 100 publications combined account for only 30% of all names. In contrast, fungal and plant names are dominated by older monographs, and 40-50% of names are included in the top 100 publications.

**Figure 12.**
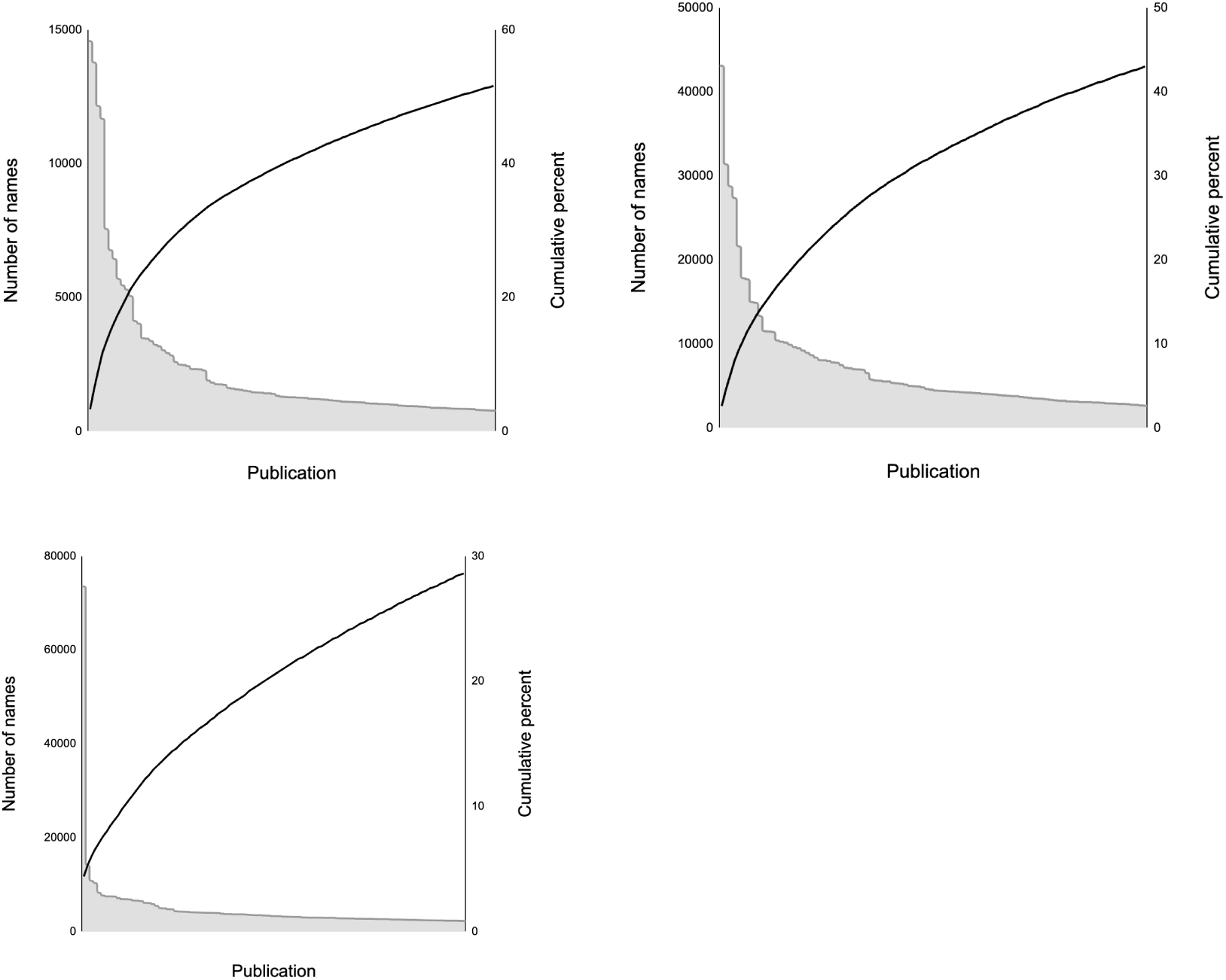
Long tails in Index Fungorum, IPNI, and ION. For each database the top 100 publications are ranked in descending order of the number of names each publishes. The left vertical axis shows the names in each publication, the right vertical axis shows the cumulative percentage of total names in the database that is added by each publication.

For fungal and plant names there is not one massively dominant publication, and the publications with the most names tend to be old monographic series from the 18th and 19th centuries.

Animal taxonomy shows a rather different pattern, with a single journal Zootaxa publishing many more names than any other. In contrast to plant and fungi names, the top n venues for publishing taxonomic names are all currently active journals.

### ChecklistBank and application

Datasets for IF, IPNI, and ION have been published to Zenodo and ChecklistBank (Table 2). These data sets are in COLDP format and comprise the subset of names from each data source that have been mapped to one or more persistent identifiers.

**Table 2.** Datasets of names mapped to persistent identifiers for the literature. Each dataset has a DOI for the data, a corresponding ChecklistBank id, and the code to generate the data is in a Github repository.

| Dataset name | DOI (all versions) | ChecklistBank ID | Github Repository |
| --- | --- | --- | --- |
| Index Fungorum | https://doi.org/10.5281/zenodo.7211134 | 129659 | https://github.com/rdmpage/index-fungorum-coldp |
| IPNI | https://doi.org/10.5281/zenodo.7208699 | 164203 | https://github.com/rdmpage/ipni-coldp |
| BioNames (ION) | https://doi.org/10.5281/zenodo.7977714 | 128415 | https://github.com/rdmpage/bionames-coldp |

Each of these can be queried using the ChecklistBank interface, or via the ChecklistBank API. As a proof of concept of what can be done with the mappings, I have released a web site called “Species Cite” https://species-cite.herokuapp.com which takes a user-supplied taxonomic name and queries the IF, IPNI, and ION datasets in ChecklistBank for a persistent identifier associated with that name. If it finds either a DOI or a Wikidata item identifier it displays those, along with a formatted citation of the paper that published the name. It also endeavours to find a PDF of that publication on the web so that the user can read more about that taxon (Fig. 13).

**Figure 13.**
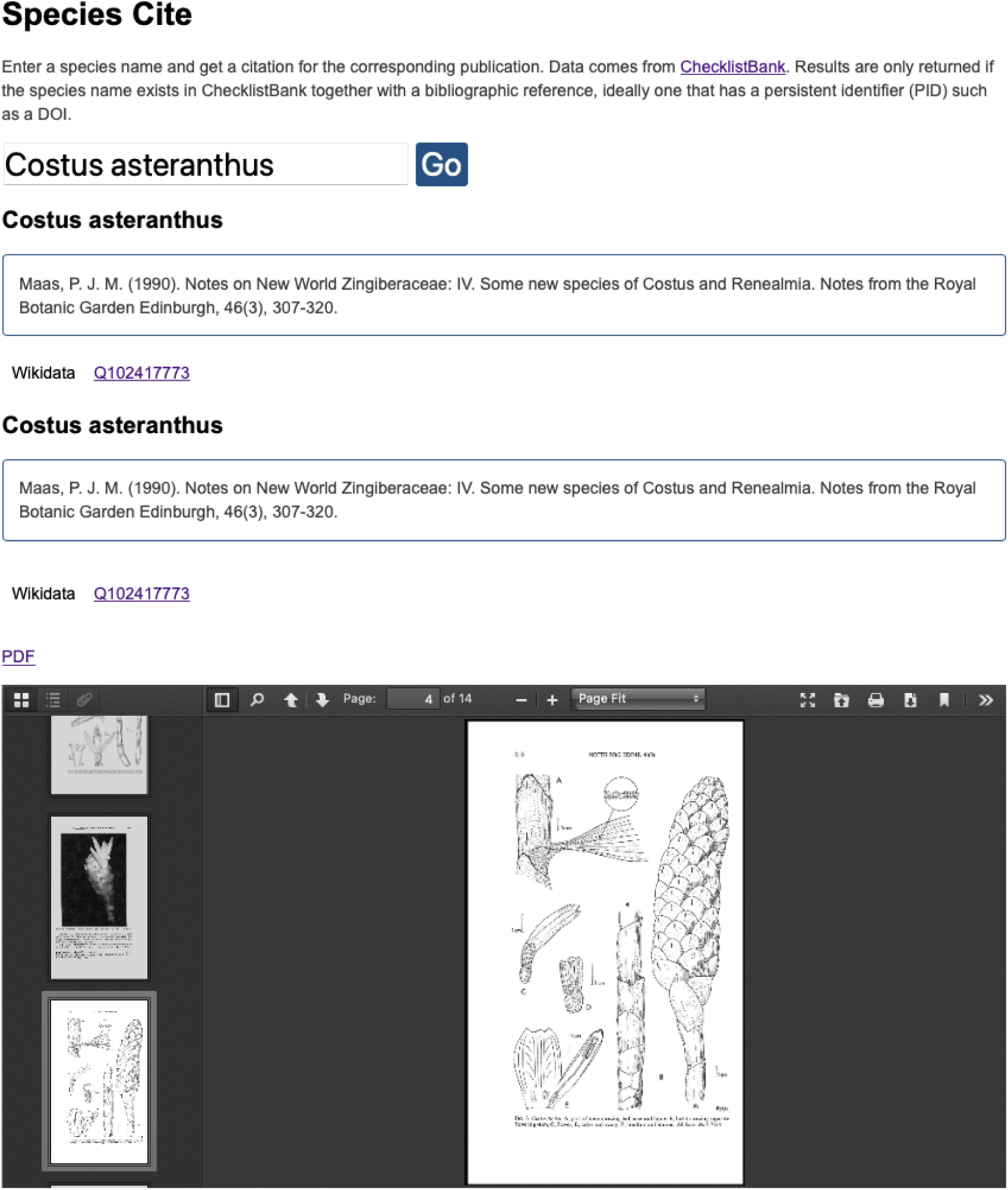
Screenshot of Species Cite showing details for a taxonomic name, with a formatted citation, links to persistent identifiers (e.g., DOI, Wikidata), and a PDF of the publication.

### Knowledge graph

The literature mappings for Index Fungorum, IPNI, and ION, together with the JSON-LD export from ORCID are available in Zenodo as RDF in N-Triples format (Table 3). The code to assemble these into a local triple store is available on Github https://github.com/rdmpage/ten-kg.

**Table 3.** The N-Triples datasets for people and taxonomic names.

| Data set name | DOI (all versions) |
| --- | --- |
| Linked data for taxonomic authors in ORCID | https://doi.org/10.5281/zenodo.7181180 |
| Index Fungorum | https://doi.org/10.5281/zenodo.7977299 |
| IPNI | https://doi.org/10.5281/zenodo.7977435 |
| ION (BioNames) | https://doi.org/10.5281/zenodo.7977556 |

The triples listed in Table 3 are minimalist in that they lack details on the taxonomic names, other than the name string and the DOI of the associated publication. Likewise, ORCID triples include a bare minimum of information about a publication, typically just the DOI and title. However, the ORCID triples also have links between people and institutions, so we can do queries such as that shown in Fig. 14 which finda authors affiliated with the Royal Botanic Gardens Edinburgh who have published plant names.

**Figure 14.**
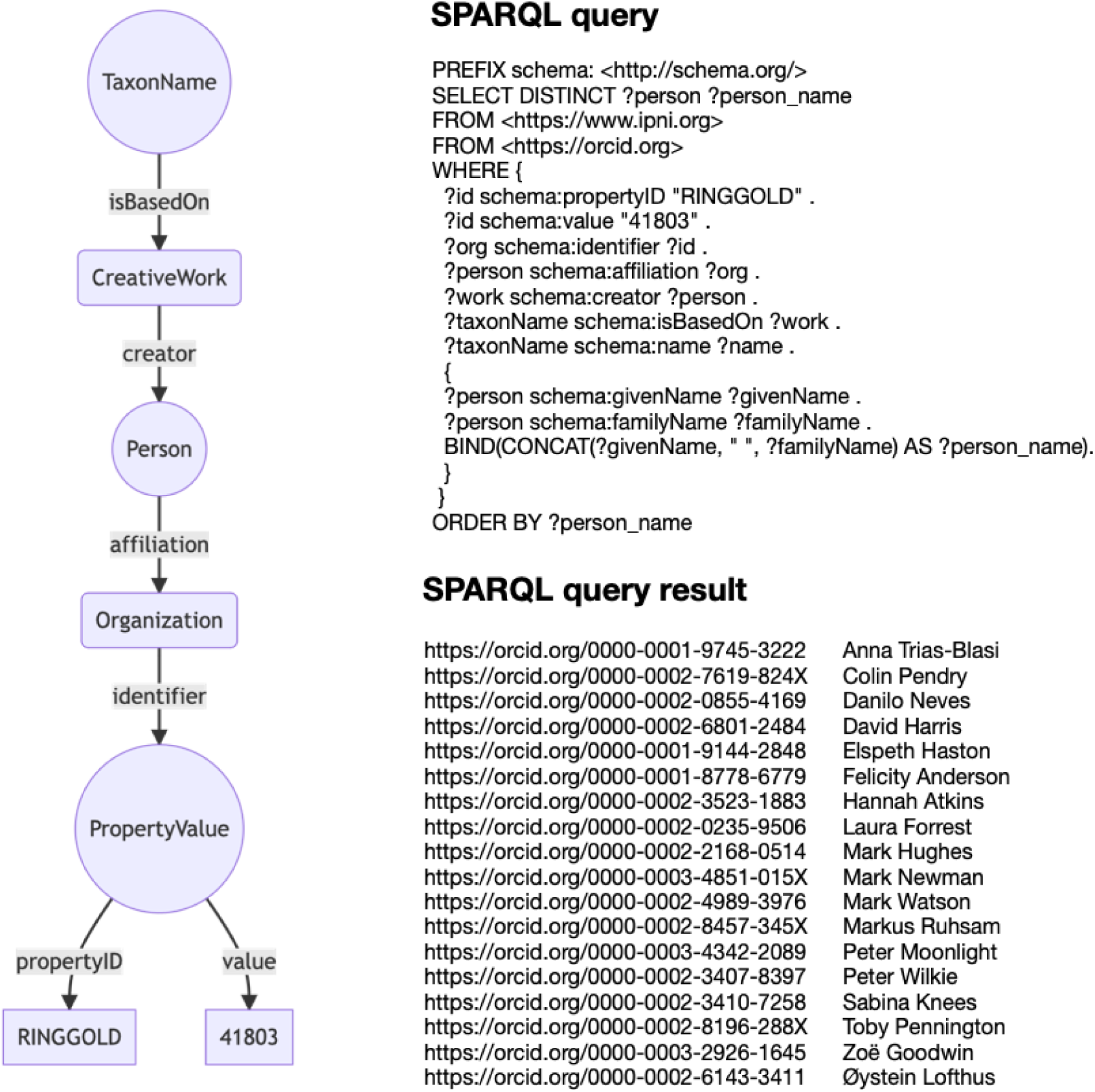
Example SPARQL query. The graph on the left connects a taxonomic name to an author of the publication of that name, and the person in turn is connected to an organisation where they worked. This graph can be represented as a SPARQL query, in this case using the RINGGOLD identifier 41803 which is for the Royal Botanic Gardens Edinburgh. The SPARQL query is restricted to the <https://www.ipni.org> namespace, so the authors listed published papers on plant names.

### Coverage

The results in Table 1 show that over a million taxonomic names have been linked to a persistent identifier for the associated publication. Overall this presents almost 20% of the total number of names in the source databases. However if we include only those names accompanied by publication information then we have approximately 36% coverage. Clearly much work remains to be done.

The long tails shown in Fig. 12 provide some insight into the problem. Even if we make rapid progress on publications that have DOIs, we quickly end up having to work with a large number of often small, obscure publications that individually contribute little, but in aggregate hold a great deal of taxonomic information. The “low hanging fruit” are quickly picked.

Readers might be surprised by the very low number of BHL page links in Table 1. This is partly because the BioNames project focuses on work-level identifiers (Fig. 3), such as DOIs. It would be possible to add BHL page links to many more of the names, using tools such as (Ower & Mozzherin, 2021), however in this project I focus on citable, work-level idenrtifiers.

### Integration

Storing the mappings in ChecklistBank enables easy use by the original taxonomic databases, or databases that reuse those taxonomic name identifiers. For example, the World Flora Online (Borsch et al., 2020) uses IPNI identifiers for many of its plant names. It could easily add detailed bibliographic data to those names by making use of the name - IPNI mapping. Likewise, UNITE species hypotheses frequently include Index Fungorum identifiers. Once taxonomic databases reuse existing persistent identifiers for names they will benefit from being able to reuse existing links to the literature.

Using persistent identifiers for the literature offers other benefits, such as increasing access to the actual publications. Identifiers such as DOIs typically resolve to a publisher’s web site, and publication itself may be behind a paywall, potentially inaccessible to a user. Tools such as Unpaywall take DOIs and discover whether freely accessible versions of that publication exist. These free versions may exist in institutional repositories, or in digital libraries such as the Biodiversity Heritage Library (Kearney, 2020).

Linking names to the literature also opens up possibilities for using summarisation techniques to generate knowledge about a taxon. For example, given the set of names applied to a taxon we could retrieve abstracts and/or full text for the associated publications, summarise that text, and develop query interfaces (e.g., chatbots) that can answer queries about the biology of that taxon.

### Missing nodes and edges in the knowledge graph

A knowledge graph consists of nodes (entities) and edges (relationships). To the extent that these are missing, the knowledge graph is incomplete. Missing nodes is an obvious weakness, we can always expand the scope and utility of a knowledge graph by adding more entities. For example, the knowledge graph described here lacks taxa (it has taxonomic names, but makes no claims about the validity of those names). One reason for this is that taxa are rarely expressed using taxon name identifiers. It is possible to retrieve JSON-LD from web pages for Catalogue of Life taxa, but the corresponding RDF lacks taxon name identifiers (the taxon names are treated as blank nodes). Hence links between taxa and names would have to be done via matching on name strings, a process that can lead to mistakes (e.g., homonyms). A significant improvement to CoL would be the use of persistent identifiers for taxonomic names provided by nomenclators such as Index Fungorum and IPNI (compare Fig. 1 and Fig. 2). Other candidate nodes are type specimens and nucleotide sequences (e.g., DNA barcodes). Most specimens currently lack persistent identifiers - there is considerable folklore about how unstable GBIF occurrence record identifiers are. Hence maintaining stable links between taxonomic names and type specimens would be a significant challenge.

### The role of citations

A potentially very useful class of missing edges are citation links between articles (the “citation graph”). Apart from the obvious, if controversial (Loizides et al., 2022; Pinto et al., n.d.), utility in developing metrics for the impact of journals, articles and researchers, and the potential for discovering related publications through co-citation, we could potentially use citation patterns as measures of the quality of taxonomic data. Taxonomists make mistakes, in the sense that they partition biodiversity up into sets (e.g., species) that subsequent research may show to be incorrect. This results in taxonomic synonyms, such as having more than one name for the same species. Solow et al. (1995) suggest that it’s not uncommon for 50% of taxonomic names to be synonyms, and note that a considerable period of time may elapse between a name being published and its eventual discovery to be a synonym. They argue that groups with few synonyms have not necessarily been blessed with very good taxonomists, rather they may suffer from neglect. If these taxa were well-studied then more synonyms would be discovered. One way to measure taxonomic activity could be citations. If the taxonomic literature of a group has received few citations, especially by other taxonomists, then this could be a clue that a group is neglected and needs more attention. Perhaps citations could be used as a proxy for taxonomic quality. At present the Catalogue of Life uses an arbitrary “star system” to rate the quality of taxonomic databases, the number of stars being self-assigned by the data provider. Citation-based measures may provide a more objective measure of the current state of knowledge of a taxonomic group.

The notion of citation could be extended to other entities, such as nucleotide sequences, such that we link DNA sequence accession numbers to the publications that cite them. Given the use of DNA sequences to identify species as well as construct phylogenies, it is likely that sequences may be cited by more than just the original publication, and indeed may link publications that don’t have any bibliographic links. That is, a subsequent paper might not cite the original publication of a DNA sequence even if it uses that sequence (R. D. Page, 2010).

### Identifier types

In this work I have focussed on "location based" identifiers such as DOIs and LSIDs. These identifiers specify a location where one can retrieve information about a digital entity, and potentially retrieve that entity itself. Location-based identifiers emphasise the persistence of resolution (for example through a centralised resolver such as https://doi.org) but typically make no guarantees that the content returned persists unchanged over time. For example, academic publishers may update the metadata for an article, but the DOI for that article remains unchanged.

It is worth noting that there is another approach to persistent identifiers, namely using cryptographic hashes of the content as the identifier (Elliott et al., 2020). This has the advantage of ensuring that the data requested hasn’t changed (which we can check by comparing the hash identifier with the hash of the data itself). Unlike DOIs and similar identifiers, there is typically no centralised mechanism to resolve hash-based identifiers. Some decentralised systems have been developed, but it is unclear if they themselves will persist. To date there are no widely used hash-based identifiers for publications.

### Next steps

This paper has described a small neighbourhood of the biodiversity knowledge graph. It is clear that there is still a considerable amount of taxonomic literature to locate and link to. The number of publications with persistent identifiers is growing, and an increasing fraction of the taxonomic literature is being retrospectively digitised. The challenge is now to ensure that this literature is made discoverable, citable, and connected to taxonomic names, thus building the bibliography of life (King et al., 2011; R. D. M. Page, 2022). We also need to develop ways to incorporate these links into existing resources, such that a visitor to a biodiversity web site is never faced with the prospect of having to Google a cryptic citation if they want to learn more about a species.

The focus of the work described here has been on bibliographic identifiers at the level of the work, that is, identifiers that are likely to be cited. This is not to discount the value in having deep links below the level of the work, such as to individual pages, or to a collection of pages (e.g., treatments). But identifiers that are cited are more likely to be the basis for new metrics of productivity (McDade et al., 2011). By including persistent identifiers for literature in taxonomic databases, we could explore mechanisms for credit for taxonomists. At present aggregators such as ChecklistBank include ORCIDs for those who contributed to curating individual databases, many of whom are taxonomists. But the bulk of the taxonomic community does not receive credit for the original work being aggregated. The use of persistent identifiers for names, publications, and people means we could start to identify those people who have contributed the most to our taxonomic knowledge. Indeed, we can envisage a case where taxonomic databases and aggregations such as the Catalogue of Life gave credit directly to the taxonomists whose data they aggregate.

